# A Bayesian framework for identifying consistent patterns of microbial abundance between body sites

**DOI:** 10.1101/673277

**Authors:** Richard Meier, Jeffrey A Thompson, Mei Chung, Naisi Zhao, Karl T. Kelsey, Dominique S. Michaud, Devin C. Koestler

## Abstract

Recent studies have found that the microbiome in both gut and mouth are associated with diseases of the gut, including cancer. If resident microbes could be found to exhibit consistent patterns between the mouth and gut, disease status could potentially be assessed non-invasively through profiling of oral samples. Currently, there exists no generally applicable method to test for such associations. Here we present a Bayesian framework to identify microbes that exhibit consistent patterns between body sites, with respect to a phenotypic variable. For a given operational taxonomic unit (OTU), a Bayesian regression model is used to obtain Markov-Chain Monte Carlo estimates of abundance among strata, calculate a correlation statistic, and conduct a formal test based on its posterior distribution. Extensive simulation studies demonstrate overall viability of the approach, and provide information on what factors affect its performance. Applying our method to a dataset containing oral and gut microbiome samples from 77 pancreatic cancer patients revealed several OTUs exhibiting consistent patterns between gut and mouth with respect to disease subtype. Our method is well powered for modest sample sizes and moderate strength of association and can be flexibly extended to other research settings using any currently established Bayesian analysis programs.

## 1 Introduction

Microbial communities inhabit virtually every part of the human body and can differ across individuals. Even within the same individual, microbial communities often change with anatomical location, Faith et al. (2013). In this context, it is not surprising that the human microbiome plays an important role in a wide range of diseases, including even life threatening conditions such as cancers. In their review, Goodman and Gardner (2018) summarize several compelling examples, such as increased *Fusobacterium* species associating with tumors in colon and *Helicobacter pylori* inducing lymphoma and gastric cancer. More recently, bacteria have been identified in pancreatic tissue in cancer patients, del Castillo et al. (2019), and have been shown to play a role in carcinogenesis in the pancreas, Pushalkar et al. (2018). Additional studies have also reported evidence that certain oral bacteria and periodontal disease associate with an increased risk in pancreatic cancer, Michaud et al. (2012); Fan et al. (2016). These findings motivate the question of whether pathologically associated microbes exist for which changes in abundance, or rate of presence, in the oral cavity correspond to changes in the abundance, or rate of presence, in gut samples. Identification of such species, exhibiting **pa**irwise **st**ratified **a**ssociation (PASTA) between two body sites, may not only allow further insight into a disease but could also provide new opportunities for treatment or detection. If the abundance of a specific microbe exhibits a consistent pattern between the two body sites with respect to disease status, then a researcher or medical professional could potentially learn about the disease in the gut by monitoring oral samples. An overview of the experimental design for testing the hypothesis of PASTA can be found in **Figure 1**. Considering that gut samples can only be acquired through invasive surgical procedures, PASTA microbes could constitute invaluable, clinical markers.

**Figure 1.**
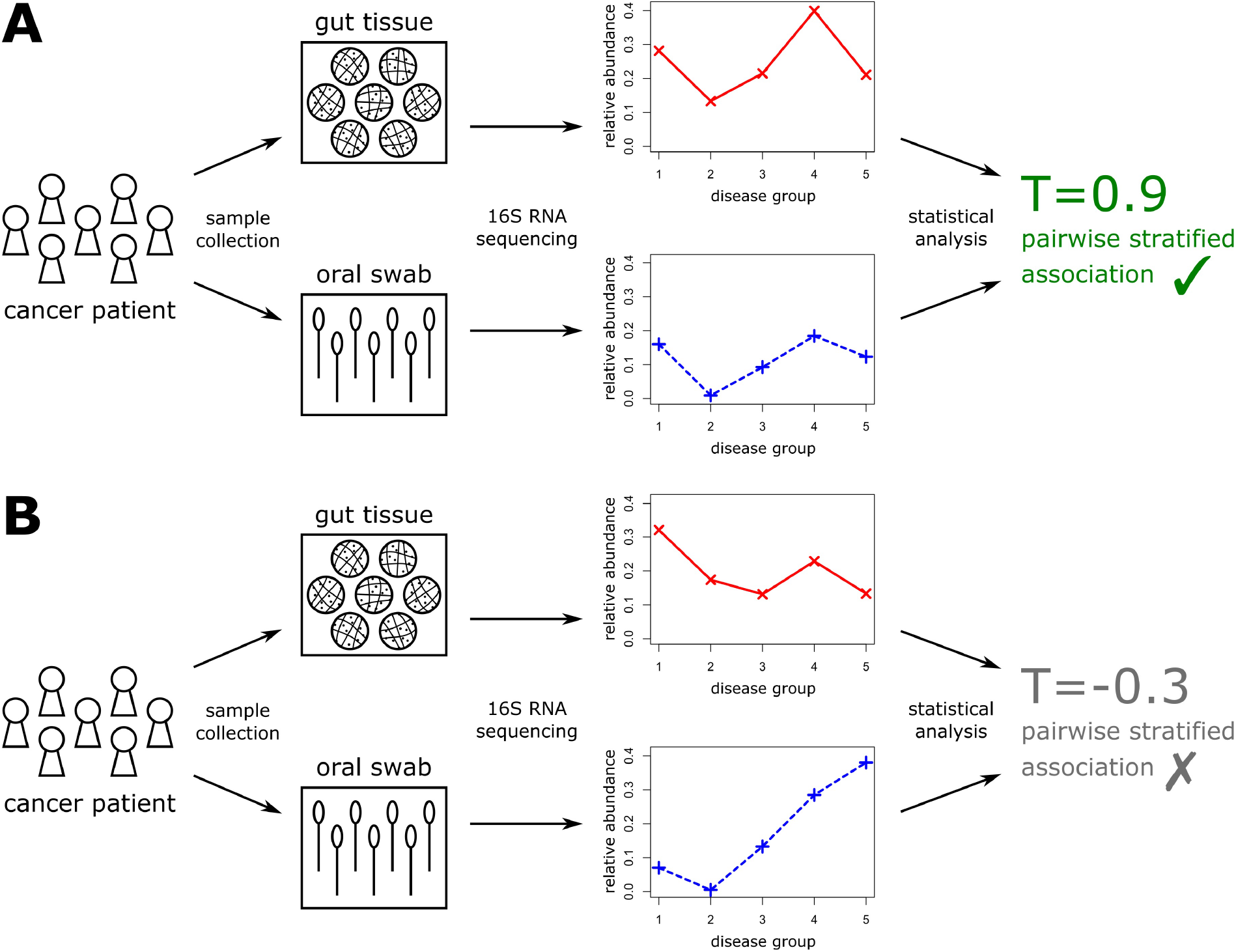
Overview of the experimental setup to test for pairwise stratified association (PASTA). Panel A depicts an example in which a consistent microbial relative abundance pattern is observed between mouth and gut, whereas panel B depicts an example in which no consistent pattern is observed.

Microbial 16S rRNA abundance data takes the form of compositional count tables in which each row represents a taxonomic unit and every column represents a sample. Unfortunately, these data are intricate with total column counts (sequencing depth and microbial yield) differing between samples, high frequency of zero values (i.e. sparsity), and the constant sum constraint problem that can create spurious associations when few rows dominate the majority of counts, Tsilimigras and Fodor (2016); Gloor et al. (2017). Due to its complexity, many different modeling strategies have been proposed for the analysis of microbial 16S rRNA abundance data. When investigating an individual microbe (or a specific group of microbes), current strategies predominantly aim to understand the relationship between abundance and selected phenotypes. Three major parametric approaches are employed by most researchers: discrete data models such as Zero-inflated Poisson or Zero-inflated Negative Binomial regression, Xia et al. (2018); Zhang et al. (2017); log-ratio Aitchison models that explicitly address the constant sum constraint by treating the ratio of abundance counts between two taxonomic units as the response, Shi et al. (2016); Tsilimigras and Fodor (2016); Gloor et al. (2017); and lastly, relative abundance models that transform counts into sample proportions and fit semicontinuous models to the data such as Zero-inflated Beta regression (ZIBR), Xia et al. (2018); Peng et al. (2016); Chen and Li (2016). Each approach can present specific advantages and limitations, where the most suitable model will depend on the circumstances of the research study. While log-ratio Aitchison models are mandatory in datasets either measuring high phylogenetic levels with few taxonomic units or exhibiting low community diversity, Tsilimigras and Fodor (2016), discrete data and relative abundance models are convenient to address sparsity in high diversity settings. To date, neither of these modeling strategies has been utilized to test for PASTA relationships and there presently exists no general testing approach that is applicable regardless of the parametric modeling strategy. Alternatively, non-parametric inter-rater strategies can be employed to test for agreement or association between body-sites. These strategies assume that there are individual raters that are presented with two different scenarios or cases, each of which they have to assign to either a category or numeric value. The methods then ask the question whether individual raters tend to make assignments that agree or associate between the two scenarios. Popular examples are Cohen’s kappa, Cohen (1960); Fleiss (1971), for categorical responses and Pearson or Spearman correlation for numeric responses, Schober et al. (2018). These methods do not necessarily require knowledge about the distribution of the response and are applicable even if there is strong disagreement or variability between individual raters. However, when applying them to microbial abundance data they do not allow for adjustment of covariates and they require availability of paired samples of the two body sites for every patient.

Here, we present an approach to test for PASTA that is applicable regardless of the data model and regardless whether all, some, or none of the samples are paired. The question of PASTA relationships with respect to body site is translated into a question of association of population parameters (such as mean relative abundance) between the two body sites. A test is then proposed based on applying a correlation statistic to parameter estimates. Testing and adjusting for paired samples is made convenient by utilizing a Bayesian modeling framework. For the purpose of illustration, this paper will focus on modeling relative abundance via a ZIBR model, though as stated before, the approach is not limited to any partiular data model. After establishing the data model and introducing the approach, viability and performance are evaluated via simulation studies and through the analysis of a biological dataset involving microbiome data collected from the gut and specific oral sites in patients with pancreatic cancer and other diseases of the foregut. Finally, strengths, limitations and opportunities for future methodological development are discussed.

## 2 Description of the Approach

### 2.1 Data Model

To briefly orient the reader, an operational taxonomic unit (OTU) can be understood as a group of closely related microbes on a given taxonomic level, for example: phylum, genus, or species. In what follows we consider abundance on two taxonomic levels: the genus and the Amplicon Sequence Variant (ASV) level, the latter representing unique biological sequences that were identified from 16S genes, Callahan et al. (2017). Note that the presented ideas hold for any taxonomic level. For a given sample, relative abundance of an OTU refers to the number of times that OTU was observed scaled by the total number of observed OTUs. It represents the proportion of times an OTU was observed in a given sample.

Let *Y*_*k*_ denote the relative abundance of a specific OTU for sample *k*. This response can be modeled as a Zero-inflated Beta distribution with probability density 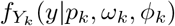. This model assumes that the case *Y*_*k*_ = 0 occurs with probability *p*_*k*_ and that given *Y*_*k*_ > 0, the response *Y*_*k*_ follows a Beta distribution with mean *ω*_*k*_ and dispersion *ϕ_k_*. For a given OTU and sample, the probability of absence *p* defines how likely it is to observe no microbe comprising that OTU within said sample. The mean non-zero relative abundance *ω* represents the mean relative abundance given that microbes comprising the OTU are actually observed. The probability density function of this distribution can be expressed as follows:

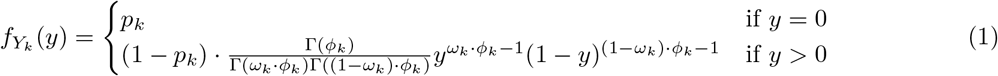

The specific ZIBR model considered here assumes that all *Y*_*k*_ are independently distributed with the common dispersion parameter *ϕ*. Let ***β***, ***δ*** denote coefficient vectors, **b**, **d** denote random effect vectors and **Q**, **R**, **W**, **X** represent design matrices. Covariates can then introduced into the model via the following link functions:

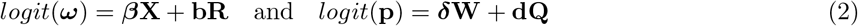

For our application, the matrices **W**, **X** are used to model the strata of body site and disease status, but can additionally be used to adjust for other, fixed covariates. On the other hand, the optional inclusion of **Q**, **R** allows to adjust for correlation structures, such as within-subject correlation when multiple samples are collected from the same patient. Posterior distributions of the coefficients were estimated via Markov chain Monte Carlo (MCMC) sampling, utilizing the following independent prior distributions for any given *t* ∈ ℕ:

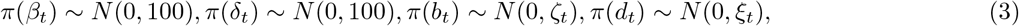

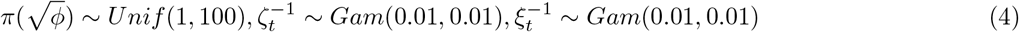

Models were fit in the software OpenBUGS (version 3.2.3 rev 1012) via the R (version 3.4.0) package “R2OpenBUGS” (version 3.2.3.2).

### 2.2 Formal Definition of Pairwise Stratified Association (PASTA)

Let *s* denote a grouping variable for which two groups are to be compared. For our purposes, this grouping variable represents body sites: *s* = 1 denotes gut and *s* = 2 denotes mouth. Let *g* denote another grouping variable with three or more distinct categories. This grouping variable will represent different types of disease status, more specifically cancer-subtype. Let *θ*_*sg*_ be a population parameter of the response for a given body site *s* and disease status *g*. The population parameter represents fundamental properties of the distribution of the response. For the here considered ZIBR model, *p* and *ω* are relevant candidates for *θ*. If PASTA holds for a given species, then either *p* or *ω* will associate between the two body sites, because they both relate to the magnitude of abundance.

We thus define: The parameter *θ* exhibits PASTA with respect to *s* and *g* if there exists an increasing function *h*(*x*) such that *θ*_1*g*_ = *h*(*θ*_2*g*_) holds for all *g* ∊ {1, 2, …, *G*}, where *G* ≥ 3. A visualization is provided in **Figure 2**.

**Figure 2.**
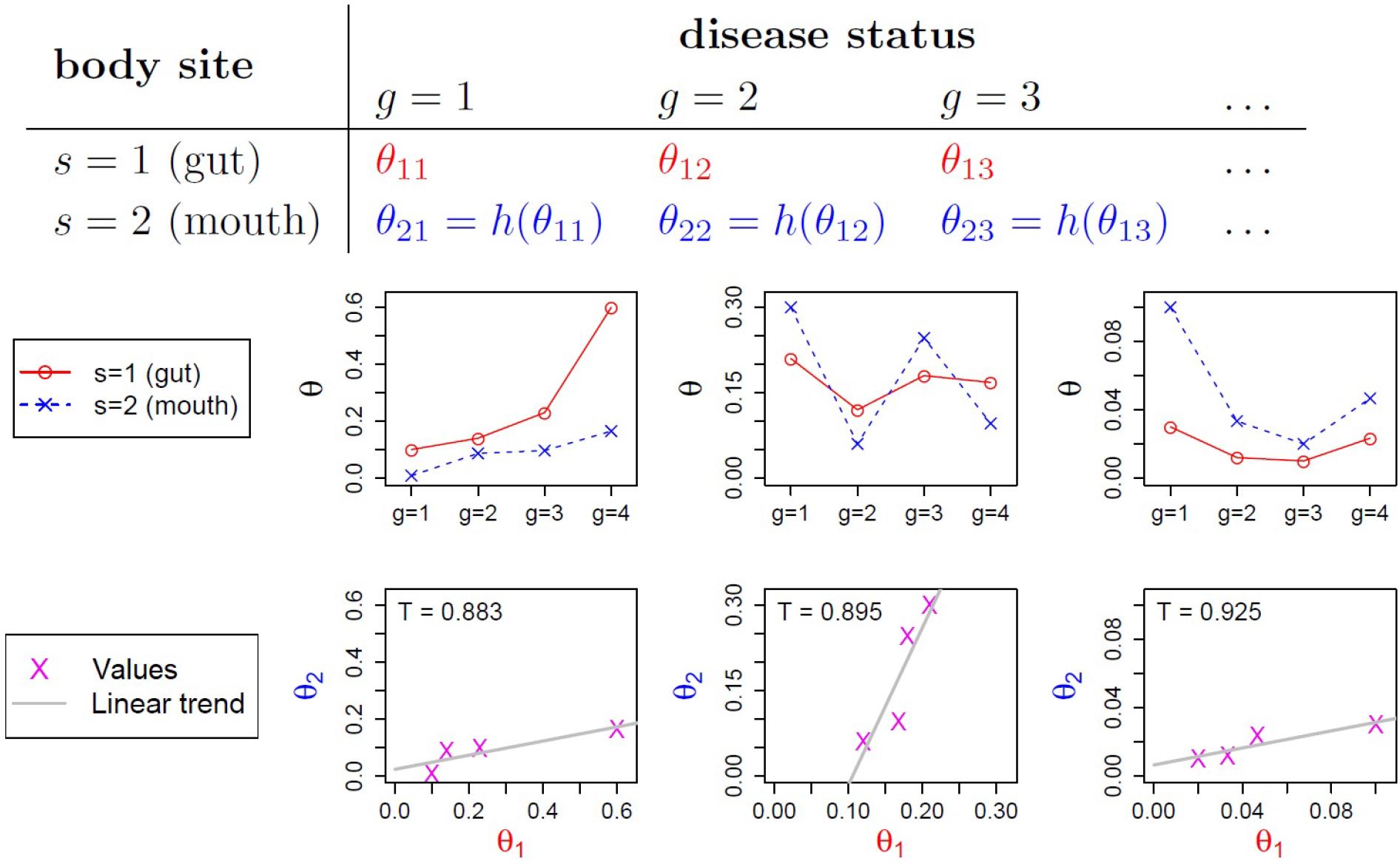
Visualization of pairwise stratified association (PASTA). Let *θ* represent a population parameter of interest, for example the mean relative abundance of a particular OTU. Each column of sub-figures below the table are examples of a PASTA relationship, i.e. of *h* being an increasing function. The first row plots parameter values of mouth and gut side-by-side and demonstrates that a variety of different scenarios are covered by this definition. In the second row, plotting parameter values of gut against parameter values of mouth reveals their association through a trend. T denotes Pearson correlation values between gut and mouth.

### 2.3 Testing for PASTA

Let *T* (**x**, **y**) ∈ [−1, 1] denote a correlation statistic between two numerical vectors **x**, **y**; for example the Pearson or Spearman correlation statistic. Also, let *T*_*θ*_ = *T* (***θ***_1_, ***θ***_2_) denote the correlation statistic for the two parameter vectors corresponding to *s* = 1 and *s* = 2. For the purpose of this paper we will assume that *θ*_1*g*_ = *h*(*θ*_2*g*_) implies *T*_*θ*_ > *t*_c_ for some −1 < *t*_*c*_ < 1. The constant *t*_*c*_ represents a meaningful degree of association. For example, a value of *t*_*c*_ = 0 would mean that any tangible degree of association is meaningful, where a value of *t*_*c*_ = 0.5 would mean that a moderate degree of association is meaningful. This definition is useful because *T*_*θ*_ scores the degree of association without explicitly having to specify the shape of *h*. Generally, the larger *T*_*θ*_ the higher the degree of association.

Based on this scoring definition of PASTA, we formulate our hypotheses in the following way:

*H*_0_: *T*_*θ*_ ≤ *t*_*c*_, i.e. ***θ***_1_ and ***θ***_2_ do NOT exhibit PASTA
*H*_1_: *T*_*θ*_ > *t*_*c*_, i.e. ***θ***_1_ and ***θ***_2_ DO exhibit PASTA

While deriving analytical solutions of the distribution of 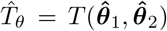 will depend on the data model and may be difficult or even impossible to obtain depending on the modeling scenario, a general testing procedure can still be derived. In the Bayesian setting, MCMC methods allow one to conveniently obtain a large sample of posterior draws of each *θ*_*sg*_, even when obtaining analytical solutions of posterior distributions is not possible. Furthermore, plugging the posterior draws of each MCMC iteration into *T* allows one to obtain posterior draws from *T*_*θ*_ itself. Let *α* denote the target credibility threshold, *H*_0_ is then rejected if the lower bound *t*_*Qα*_ of the one-sided credible interval of *T*_*θ*_|**Y** exceeds *t*_*c*_. This is equivalent to rejecting *H*_0_ if the posterior probability of no association exceeds *α*, i.e. *P r*(*T*_*θ*_|***Y*** < *t*_*c*_) < *α*. In detail, the step by step process for testing *H*_0_ is as follows:

1. Specify a likelihood for the response data **Y** and prior distributions for the parameters ***θ***
2. Utilize a MCMC sampling scheme to draw a large number of samples from the posterior distributions of the parameters ***θ*** | **Y**. One draw from the Markov Chain contains a unique draw for each *θ*_*sg*_.
3. Calculate 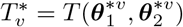 where ***θ***^∗*υ*^ denotes the *υ*^*th*^ MCMC draw. Then **T**\* is a large sample of the posterior distribution *T*_*θ*_|**Y**.
4. Calculate the *α* · 100% sample quantile *t*_*Qα*_ of **T**\*. If the Markov Chain is sufficiently long, the sample quantiles of **T**\* will closely approximate the quantiles of the true posterior distribution. The value *t*_*Qα*_ is thus the lower bound of the (1 − *α*) · 100% one-sided credible interval of *T*_*θ*_|**Y**.
5. Reject *H*_0_ if the lower bound *t*_*Qα*_ is larger than *t*_*c*_.

This process is generally applicable regardless of the data model or the parameter being tested, as long as each *θ*_*sg*_ can be estimated without constraining them to a parameter space that implies PASTA.

## 3 Results

### 3.1 Simulation Studies

Performance of our proposed approach was first evaluated using series of simulation studies. In an attempt to obtain sampling distributions of parameters that would approximate biological distributions, unstratified ZIBR models were fit to each OTU in the pancreatic cancer dataset (see Methods for details of this dataset). Unstratified parameter estimates were then used to obtain smooth sampling distributions of *ω*, *p*, *ϕ*. Finally, these sampling distributions were used to generate many pseudo-datasets satisfying *H*_1_ and performance was evaluated when applying the previously described testing approach to the simulated dataset.

Sampling distributions for parameters were similar for both genus and ASV level. However, for *ω*, the mean non-zero relative abundance, distributions tended to be slightly further concentrated toward 0.0 on the ASV level as compared to the genus level. Further, distributions of *p* tended to be slightly more concentrated toward 1.0 on the ASV level as compared to the genus level. In both cases a linear relationship was observed between log *ω* and log *ϕ* which was ultimately used to sample *ϕ* conditionally on *ω* (**Figure 3**).

**Figure 3.**
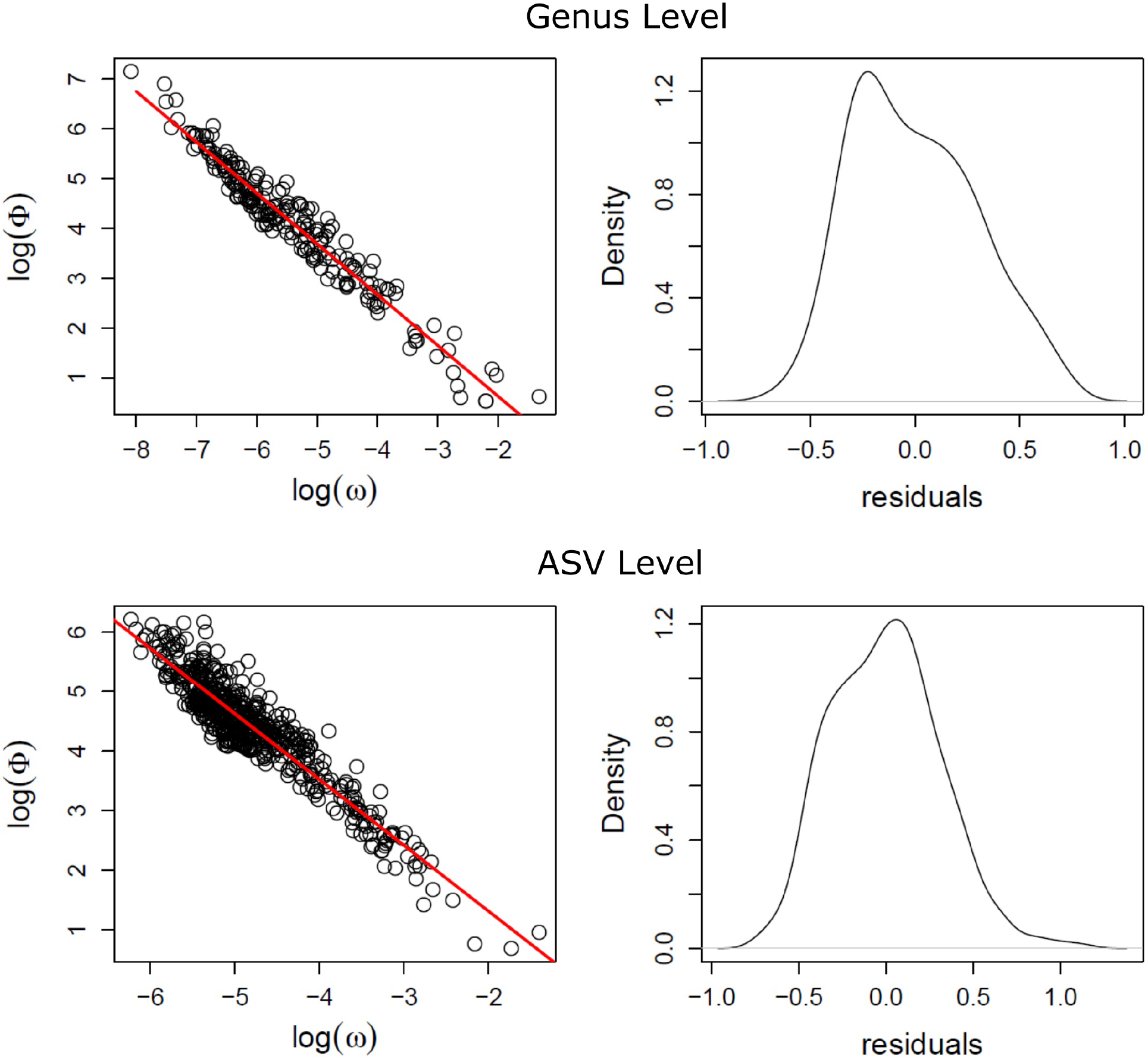
Observed relationships between marginal distributions of *ω* and *ϕ* estimated from the pancreatic cancer dataset. For both the genus and the ASV level, parameters were estimated marginally for each OTU across all observation without any stratification. When plotting marginal parameter estimates of *ω* and *ϕ* a linear relationship can be observed on the log scale. This relationship was utilized to sample *ϕ* conditionally on *ω* in the simulation studies.

In summary, the following sampling distributions were obtained:

Genus: *p* ~ *Beta*(1.67, 0.4); *ω* ~ *Beta*(0.63, 53.27); log *ϕ*| min_*sg*_{log *ω*_*sg*_}) ~ *N* (−1.02 min_*sg*_{log *ω_sg_*} − 1.41, 0.3^2^)
ASV: *p* ~ *Beta*(7.35, 0.49); *ω* ~ *Beta*(1.46, 121.12); log *ϕ*| min_*sg*_{log *ω*_*sg*_}) ~ *N* (−1.10 min_*sg*_{log *ω*_*sg*_} − 0.89, 0.31^2^)

As expected, simulations of biological data revealed that analyses on the genus level were overall more powerful than on the ASV level, regardless of which population parameter was investigated (**Figure 4**). Assuming *t_c_* = 0, four disease status groups, 95% credible intervals and utilizing Pearson correlation, the highest power was achieved when testing PASTA of *ω*. Under a moderate degree of association of *T*_*θ*_ = 0.537 a target power of 0.8 was reached for 5 samples per stratum on the genus level and 15 samples per stratum on the ASV level. Type 1 error rates appeared adequately calibrated to the 5% significance level ranging from 0.03 to 0.056 on the genus level and from 0.032 to 0.06 on the ASV level. Performance for the probability of absence *p* was substantially worse. Under a high degree of association of *T*_*θ*_ = 0.834 a target power of 0.8 was reached for 40 samples per stratum on the genus level. On the ASV level, utilizing as many as 80 samples per stratum resulted in a power of only 0.59 for the same *T*_*θ*_. Type 1 error rates also appeared mostly calibrated in this scenario, but showed deflatation for smaller sample sizes, assuming a value of 0.027 on the genus level and 0.009 on the ASV level.

**Figure 4.**
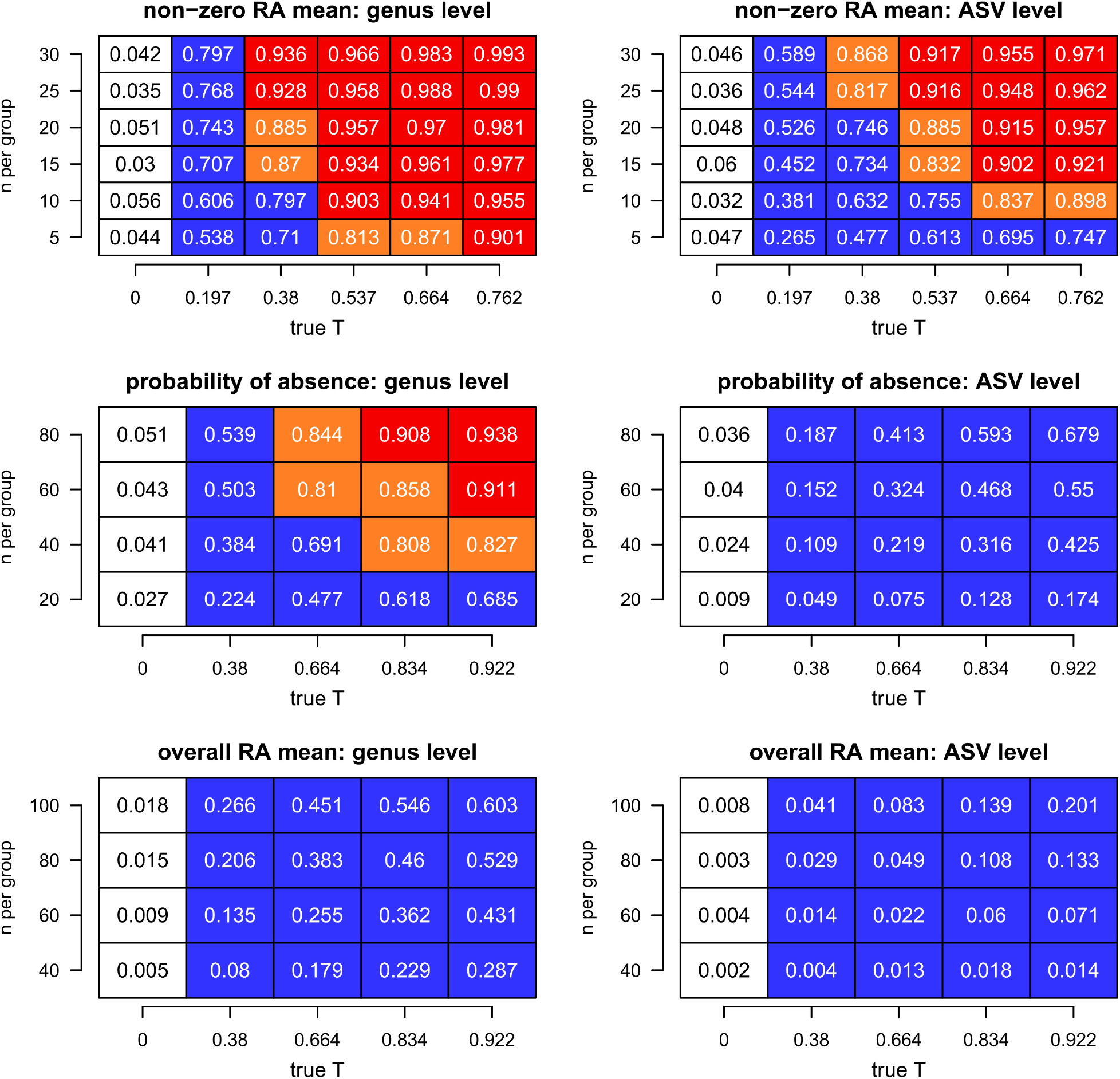
Results of the simulation studies. Power plots are displayed for testing PASTA of various population parameters with *t*_*c*_ = 0 at both ASV and genus level. The term “n per group” refers to the number of samples available in each of the eight sub-group combinations resulting from two body sites and four different levels of disease status. *H*_0_ was rejected if *Pr*(*T*_*θ*_|**Y** < 0) < 0.05. Type 1 error rates are displayed in white colored boxes with black fonts. Power values less than 0.8 are colored blue, values larger than 0.9 are colored red and values between 0.8 and 0.9 are colored orange. Genus level pseudo data generally has higher statistical power than the ASV level. High performance is achieved by the non-zero mean *ω*, while an increased sample size is required for the probability of absence *p*. Tests of the overall mean *μ* result in low performance, when only mildly constraining sparsity.

Testing PASTA of the overall mean *μ* = *ω*(1 − *p*) was also investigated. While improving with increasing size of effect and sample size, the power for this parameter was lower than when considering *ω, p* individually. Even when considering the large degree of association *T*_*θ*_ = 0.834 and using 100 samples per stratum, the genus level scenario achieved a power of only 0.546. Notably, type 1 error rates were consistently deflated, ranging from 0.005 to 0.018 on the genus level and 0.002 to 0.008 on the ASV level. Type 1 error rates were deflated across all simulated scenarios, reaching values of less than or equal 0.018 or less.

Discrepancies in performance were found to be directly related to precision of parameter estimates. When plotting the posterior means of *T*_*θ*_ against their true simulated values across various simulation runs, the variation around the identity line consistently increased from *ω* to *p*, aswell as from genus to ASV level (**Figure 5**). Analogously, posterior distributions of *T*_*θ*_ were found to on average become more diffuse and more biased towards 0, when moving from *ω* to *p* or from genus to ASV level. When performing simulation runs of a scenario with low relative precision, in which *p* was sampled from a Uniform(0.85,0.95) distribution, the posterior distribution of *T*_*θ*_ was on average almost perfectly centered at zero and highly diffuse.

**Figure 5.**
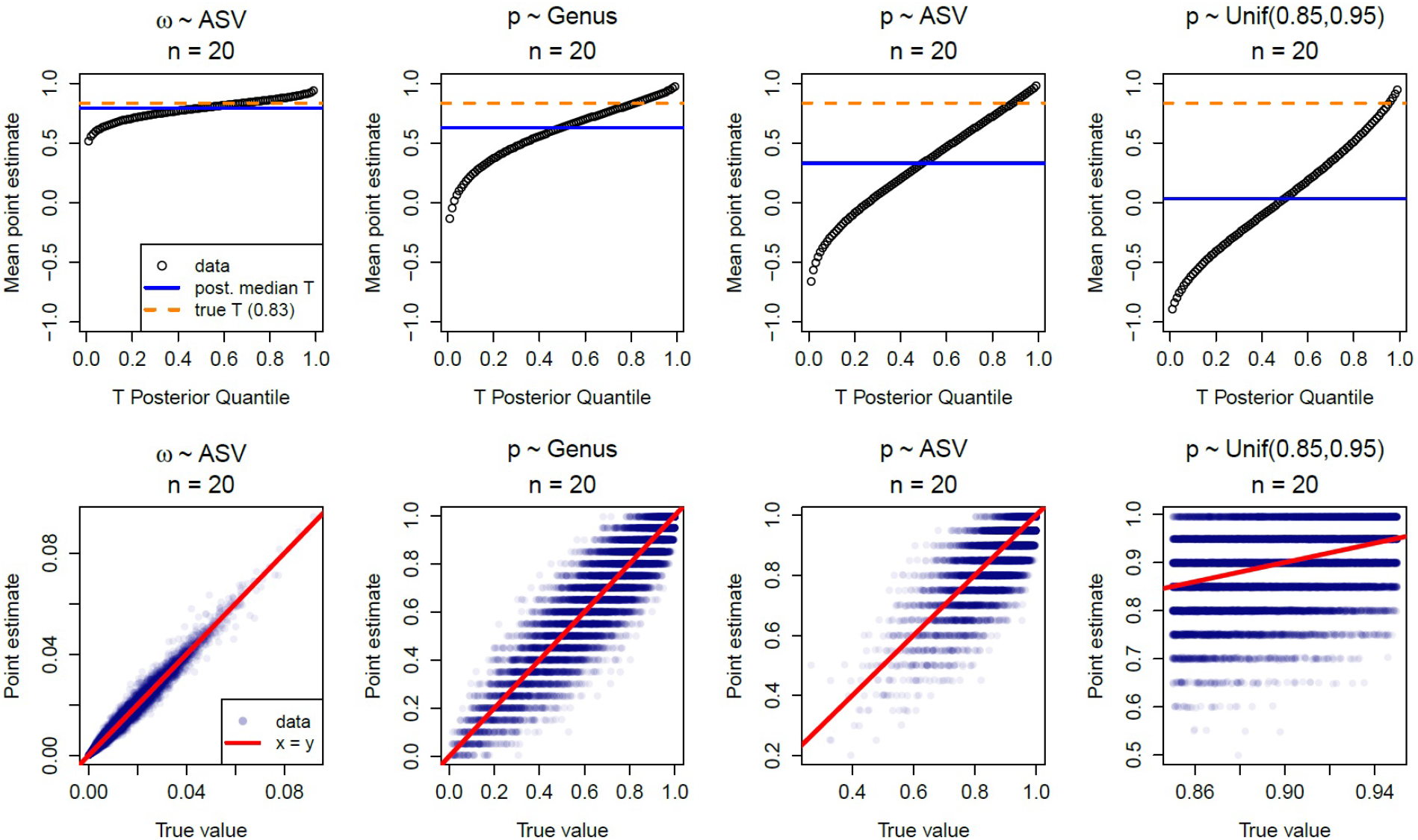
Effects of the relative precision of parameter estimates on the posterior distribution of *T*_*θ*_. The first row shows the average point estimate of posterior quantiles of *T*_*θ*_ across simulation runs for various simulation scenarios. The second row shows the associated plots of the parameters’ posterior means versus their true values across simulation runs. As the relative precision of parameter estimates decreases, the posterior distribution of *T*_*θ*_ becomes more diffuse and more biased towards 0.

Poor power when testing *μ* was also found to be related to two additional factors. Detailed results of simulations accounting for these factors are displayed in **Additional File 1**. The deflated type 1 error rates when utilizing 95% credible intervals, lead to overly conservative tests that negatively affected power. Calibrating type 1 errors to 5% by adjusting lower bounds of the credible intervals of *T*_*θ*_ for each considered sample size, lead to a consistent improvement in power, reaching a value of 0.73 for *T*_*θ*_ = 0.922 and 80 samples per stratum on the genus level. The second factor that affected performance was the employed liberal three sub-strata rule, which allowed up to three strata to exhibit exclusively zeroes. Since high rates of absence were simulated, this case often naturally occurred leading to the three respective posterior estimates being imputed with the vague prior distribution, which is very imprecise. A follow-up simulation restricting all strata to have at least one non-zero observation, lead to a consistent increase in power, reaching a value of 0.82 for *T*_*θ*_ = 0.922 and 80 samples per stratum on the genus level. In both settings, overall performance for testing *μ* was consistently lower than for testing *p* regardless of taxonomic levels. When both calibrating type 1 error and restricting non-zero observations at the same time, power increased further but was not consistently better than for testing *p*.

### 3.2 Applying the Approach to Biological Data

The 10th revision of the International Statistical Classification of Diseases and Related Health Problems (ICD-10), is a systematic classification of medical conditions provided by the World Health organization. Disease status information in this dataset was available via ICD-10 codes for each subject. The clinical pancreatic cancer dataset contained four predominant cancer-types: C24.x, C25.x, K86.2 and other, where “.x” denotes a further sub-type that could differ by subject and “other” refers to pancreatic cancer in various other categories or other diseases of the foregut. OTUs exhibiting significant PASTA with respect to disease status were successfully identified for both genus and ASV level in this dataset. For analysis we considered coding cancer sub-type in two ways: four group coding as described above and three group coding, which collapsed “K86.2” and “other” into one group. On the genus level, when coding disease status into four groups three genera exhibiting PASTA between mouth and gut were identified: *Fusobacterium*, *Haemophilus* and *Veillonella* (**Table 1**). After substratifying oral sites into saliva, tongue, buccal and gum, these association were found to be preserved for some of the site pairs: *Fusobacterium* also exhibited PASTA between gut an saliva sites; *Haemophilus* also exhibited PASTA between gut and gum, as well as gut and tongue; *Veillonella* also exhibited PASTA between gut and gum. Several genera also exhibited PASTA between individual mouth sites (**Table 2**). Two genera exhibited PASTA between four pairs of mouth sites: *Fusobacterium* and *Actinomyces*. Three genera exhibited PASTA between two pairs of mouth sites: *Atopobium*, *Haemophilus* and *Prevotella*. Six genera exhibited PASTA in only one pair of mouth sites.

**Table 1.** Genus level OTUs showing evidence of PASTA between gut and mouth sites when dividing ICD10 code into four groups. For a given genus, a parameter is included in this table if it was either marginally significant or significant when *T* was either Pearson or Spearman correlation. For a given population parameter *θ*, marginal significance (*Pr*(*T* | **Y** < 0) < 0.1) is denoted by *θ^..^* and significance (*Pr*(*T* | **Y** < 0) < 0.05) is denoted by *θ*\*. Three parameters were investigated: *μ, ω, p*. Due to low power in this exploratory setting multiple testing was not adjusted for.

| genus | gut &<br>mouth (all) | gut &<br>buccal | gut &<br>gum | gut &<br>saliva | gut &<br>tongue |
| --- | --- | --- | --- | --- | --- |
| *Fusobacterium* | $\mu^{\cdot\cdot}, p^{\cdot\cdot}$ | - | - | $\mu^{\cdot\cdot}$ | - |
| *Haemophilus* | $p^*$ | - | $\mu^*, p^*$ | - | $p^{\cdot\cdot}$ |
| *TM7-G1* | - | - | - | - | $p^{\cdot\cdot}$ |
| *Veillonella* | $p^{\cdot\cdot}$ | - | $p^{\cdot\cdot}$ | - | - |

**Table 2.**
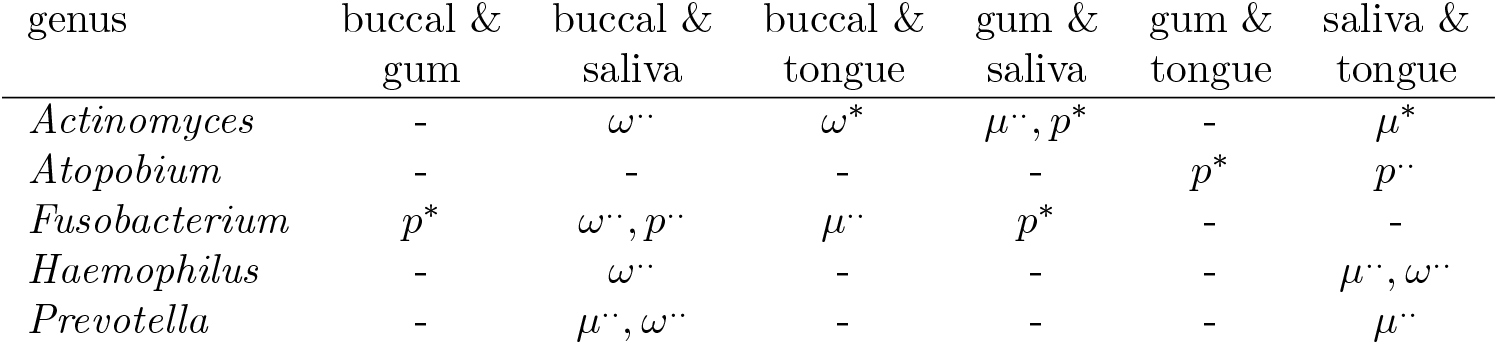
Genus level OTUs showing evidence of PASTA between mouth sites when dividing ICD10 code into four groups. For a given genus, a parameter is included in this table if it was either marginally significant or significant when *T* was either Pearson or Spearman correlation. For a given population parameter *θ*, marginal significance (*Pr*(*T* | **Y** < 0) < 0.1) is denoted by *θ*^..^ and significance (*Pr*(*T* | **Y** < 0) < 0.05) is denoted by *θ*\*. Three parameters were investigated: *μ, ω, p*. Due to low power in this exploratory setting multiple testing was not adjusted for. Six OTUs showing association for only one pair of mouth sites are not shown in this table.

| genus | buccal &<br>gum | buccal &<br>saliva | buccal &<br>tongue | gum &<br>saliva | gum &<br>tongue | saliva &<br>tongue |
| --- | --- | --- | --- | --- | --- | --- |
| *Actinomyces* | - | $\omega^{\cdot\cdot}$ | $\omega^*$ | $\mu^{\cdot\cdot}, p^*$ | - | $\mu^*$ |
| *Atopobium* | - | - | - | - | $p^*$ | $p^{\cdot\cdot}$ |
| *Fusobacterium* | $p^*$ | $\omega^{\cdot\cdot}, p^{\cdot\cdot}$ | $\mu^{\cdot\cdot}$ | $p^*$ | - | - |
| *Haemophilus* | - | $\omega^{\cdot\cdot}$ | - | - | - | $\mu^{\cdot\cdot}, \omega^{\cdot\cdot}$ |
| *Prevotella* | - | $\mu^{\cdot\cdot}, \omega^{\cdot\cdot}$ | - | - | - | $\mu^{\cdot\cdot}$ |

On the ASV level, two ASVs exhibited PASTA with respect to *p* between mouth and gut when disease status was coded into four groups. When coding disease status into three groups, the same two ASVs as before and three additional ASVs exhibited PASTA with respect to *p* between mouth and gut. Notably, among these additional ASVs was a candidate belonging to the *Fusobacterium* genus. Further details of the ASV level analysis are discussed in Chung et al. (2019).

## 4 Discussion

The methodology presented in this publication successfully establishes a general framework to test for pairwise stratified association (PASTA) in microbial abundance or relative abundance. The approach first estimates posterior distributions of population parameters ***θ***|**Y** within the strata of body site and disease status and subsequently calculates a correlation statistic *T*_*θ*_ between body sites, which scores their degree of association. This allows researchers to identify individual microbes or groups of microbial species that show consistent abundance patterns between different body sites with respect to the disease status of patients or any other relevant categorical grouping variable.

It has to be noted that while many possible *T* may be adequate to detect a wide variety of increasing relationships *h*, the choice of *T* can favour certain shapes of *h*. If the Pearson correlation is employed, linear relationships will achieve higher scores than rapid, exponential growth relationships, since it measures the degree of linear association. In this case, overly large values of *t_c_* (for example *t*_*c*_ = 0.8) may lead to falsely rejecting non-linear, increasing relationships. While rank-correlation measures such as the Spearman correlation may be more generally applicable, they may also be less powerful, especially when few groups are considered (*g* < 5). In these cases with a small number of groups, the discrete nature of the rank-correlation statistic is more pronounced. When utilizing Spearman correlation it is helpful to keep in mind that *T*_*θ*_ can only assume 4 discrete values when *g* = 3, 11 discrete values when *g* = 4 and 21 possible values when *g* = 5. Care should also be exercised when interpreting significant associations. The test for PASTA is concerned with trend, agreement or association between *s* = 1 and *s* = 2 after stratification according to *g*, but does not at all provide information on whether the effect of site *s* or disease status *g* is biologically or clinically significant. To the contrary, it assumes that both grouping variables are inherently meaningful objects of the research hypothesis. For example, if there is no significant effect of body site (i.e. abundance is the same between mouth and gut), but abundance differs by disease status, the test statistic will likely score a high degree of association, because what is going on in one site is still associated with what is going on in the other site and this is an inherently meaningful relationship to us. However, the contrary where effect of body site is significant (i.e. abundance is different between mouth and gut) but effect of disease status is not, will not necessarily lead to a significant score of association. Scenarios are possible in which there are small effects of body site and disease status, where none are strong enough to reach statistical significance, yet the test for association may still be overall significant, as long as the trend across strata is pronounced enough. To understand the specific nature of an identified PASTA relationship it can be useful to plot credible intervals of parmaeter estimates *θ_sg_* side-by-side (**Additional File 2**) or to perform statistical follow-up tests investigating the effects of *s* and *g*.

Results of the simulation studies reveal that the testing procedure is able to successfully identify PASTA patterns. The decreased performance on the ASV level can be attributed to decreased signal intensities and the overall increase in sparsity of non-zero observations. The substantial drop in performance when investigating PASTA of *p* was demonstrated to be a result of overall lower precision in estimation, compared to *ω*. Since probabilities of absence are generally high and concentrated towards 1.0 across OTUs and strata, the differences between them are often small. In this scenario, to be able to reliably quantify differences and assess trends with adequate precision, larger sample sizes are required. This problem is thus a limitation of the zero-inflated data and not the testing approach itself.

Investigating the overall mean *μ*, may not always be viable when utilizing the ZIBR model. Since it’s estimation is based on estimates of both *p* and *ω*, it’s estimates are subject to more sources of variation, resulting in poorer precision and lower power. Our simulation suggests that if the properties of the population that is to be analyzed are well known, adjusting the quantile *t_Qα_* to calibrate type 1 error rates is a viable strategy to improve performance. If this was not the case and a researcher was convinced that inference based on *μ* was more biologically meaningful than considering the individual components *p* and *ω*, alternative models may be considered. For relative abundance data an adequate choice may be the marginalized ZIBR model as proposed by Chai et al. (2018) which directly estimates *μ* as a function of covariates. These estimates could then be used analogously to test for PASTA relationships using the here proposed approach.

The fact that in the pancreatic cancer patient dataset OTUs can be identified that show associations between mouth and gut, as well as between individual oral sites suggests that they may be promising candidates for potential biomarkers. Among these were *Fusobacterium* and *Haemophilus*, both oral bacteria recently found to distinguish pancreatic head carcinoma patients from healthy subjects, Lu et al. (2019). Also, species belonging to the genera *Fusobacterium* and *Prevotella* (even though the latter was only found to show association between mouth and gut) have been shown to associate with periodontal disease, Chiranjeevi et al. (2014); Chen et al. (2018). These results lend further credence to the disease related connection between microbial abundance in mouth and gut and suggests that our method leads to conclusions consistent with the literature. More future research will be needed to validate these findings.

The simulation studies also confirmed that tests of PASTA applied to the pancreatic cancer patient dataset are likely underpowered due to the limited sample size. It should be noted that many OTUs could not be tested due to too high zero-inflation and thus insufficient signal. These two factors likely explain why relatively few candidates were identified when conducting the tests. Future studies may consider larger sample sizes or aim to improve the yield of observed counts in each sample to alleviate this issue. Our results suggest that differences in the extend of zero-inflation between groups may be generally hard to detect for small to medium sized studies when more granular phylogenetic levels are targeted.

## 5 Conclusions

In conclusion, the performed simulation studies demonstrate the viability of the approach in the context of ZIBR models and suggest that modest sample sizes can achieve adequate power for moderate degree of association. The simulations also highlight potential lack of power for low-level phylogeny data or when more complex functions of population parameters are considered. When analyzing a biological dataset consisting of pancreatic cancer patients the approach is able to identify microbes that exhibit PASTA patterns and are consistent with independent findings of current research studies. The generality of this approach allows it to be extended to other data models and research settings, ensuring that it can be useful for researchers interested in stratified associations in the microbiome world and beyond.

## 6 Methods

### 6.1 Pancreatic Cancer Patient Dataset

In order to evaluate validity of the approach in the context of microbiome data, analyses were performed based on a biological 16S rRNA sequencing dataset first published in del Castillo et al. (2019). This dataset contained samples of various gut and oral sites from 77 patients with pancreatic cancer with age range 31 to 86 years. Sequencing was performed utilizing the Illumina MiSeq System and alignments were performed using BLASTN against a reference library combining sequences from HOMD (version 14.5), Greengenes Gold and the NCBI 16S rRNA reference sequence set. OTU counts were obtained utilizing the QIIME (Quantitative Insights Into Microbial Ecology 22) software package version 1.9.1, while the unique Amplicon Sequence Variant (ASV) counts were calculated using the QIIME2 software package release 2018.4. The former was used to obtain taxonomic genus level counts, whereas the latter was used to obtain rarefied ASV level information, calculated based on sequencing data rarefied at a sampling depth of 1200. Both genus level and ASV level counts were considered for analysis. Before fitting statistical models to the data, relative abundance values of less than 0.01 were treated as noise and set to 0.

The dataset was used to both guide simulation studies (described in the next section) and to deploy models to identify potential microbes that may exhibit a PASTA pattern.

### 6.2 Simulation Studies

Before simulations were performed, an empirical approach was pursued in order to obtain sampling distributions of the parameters *p*, *ω*, *ϕ* that would be representative of biological microbiome data. First, a marginal, unstratified ZIBR model was fit to the pancreatic cancer dataset that assumed all samples of relative abundance for a given OTU originated from the same distribution. These model fits yielded a single estimate of *p*, *ω* and *ϕ* for each OTU. These estimates were then assumed to be representative of or approximate the true distribution of parameters in biological data. In the next step, the estimates were used to obtain smooth probability distributions that parameters could be sampled from during the simulation studies. For both *p* and *ω*, individual Beta distribution models were fit to the marginal estimates in order to obtain their smooth sampling distributions. On the other hand, log *ϕ* was sampled via a Normal distribution through an observed linear relationship between log *ϕ* and log *ω* that was present on the ASV level and the genus level. More specifically, since our models assumed fixed dispersion among all groups, dispersion was sampled from (log *ϕ* | min_*sg*_ {log *ω*_*sg*_}) ~ *N* (*a* min_*sg*_ {log *ω*_*sg*_} + *b*, *σ*^2^), where *a, b, σ*^2^ differed between genus and ASV level. After the smooth sampling distributions were obtained, the performance of PASTA tests was evaluated via simulations. Let *t* denote a target, fixed degree of association, *n* denote the number of observations in each stratum (*s, g*) and *t*_*c*_ = 0 denote the tested degree of association. A single simulation run was carried out by first randomly drawing all *θ*_*sg*_ parameters from the representative sampling distributions, until |*T*_*θ*_ − *t*| < 0.001 was satisfied. This process yields parameters that are both representative and that also exhibit a target degree of association (within a small error margin). Next, the drawn parameters satisfying this condition were plugged into the likelihood of the ZIBR data model, which was in turn used to draw a random sample of relative abundance values. This simulated pseudo-data was then used to fit the Bayesian ZIBR model and conduct our hypothesis test. Each considered scenario was simulated 1000 times and statistical power for given *t*, *n* and *t*_*c*_ was then estimated as the proportion of times *H*_0_ (i.e. *T*_*θ*_ ≤ 0) was rejected. We specifically considered Pearson correlation as choice for *T* (*x, y*) in this simulation.

An additional restriction was put in place for sampling pseudo-data in order to prevent rare cases of sparse datasets with insufficient signal to perform the analysis. If a generated pseudo-dataset contained more than three sub-strata (*s, g*) in which all observations exhibit a response value of either all 0 or all 1, then it was rejected and a new pseudo-dataset was sampled.

### 6.3 Model fitting

Let *sgi* denote the *i*^*th*^ observation in the stratum for site *s* and disease status *g*. Let *X*_*sgi*_ be the log of total sample abundance for sample *sgi* and let 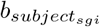 denote the random intercept for the subject that sample *sgi* originated from. The three different models that were utilized in this study are shown below:

Model A: *logit*(*ω*_*sg*_) = *β*_*sg*_ & *logit*(*p*_*sg*_) = *β*_*sg*_
Model B: 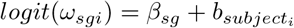 & 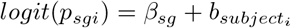
Model C: 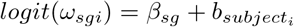 & 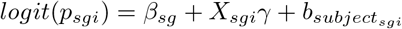

Model A was utilized in the simulation studies. Model B was utilized for fitting ASV level data, while Model C was utilized for fitting genus level data. This choice was made because scaling OTU counts to relative abundance will only make non-zero relative abundance comparable between samples, but not the rate of absence. This is due to the fact that, even if the true probability of absence *p* for a specific OTU is very high, if more microbes are overall observed in sample 1 than in sample 2, then the probability of observing none of the microbes belonging to the target OTU in sample 1 is much lower than in sample 2. For example, if a total of 1,000,000 microbes live in a body site and 100 of them belong to the genus *Prevotella*, then if we randomly extract 1,000 microbes from this body site with our sample, we would expect to only rarely find one of these 100 microbes in our sample. However, if our sample randomly extracts 100,000 microbes from the body site, it would be rare to find none of the 100 microbes in it that belong to the genus *Prevotella*. So since the genus level data was not rarefied, the total sample abundance differed between samples and an adjustment was necessary, whereas the ASV level data was rarefied and did not require adjustment for total sample abundance.

In order to achieve potentially better convergence behaviour and to simplify and speed up the model fitting, the logistic regression component of the model was fit independently of the Beta regression component, in all cases. The resulting posterior chains of *p* and *ω* were then used to calculate the posterior chain of *ω*. This approach is justified under the assumption that *p* and *ω* are independent after adjusting for covariates, but may be inadequate when there are confounders affecting both parameters not accounted for in the model.

## Declarations

### Ethics approval and consent to participate

The study was approved by Lifespan’s Research Protection Office for recruitment at RIH, as well as the Institutional Review Boards for Human Subjects Research at Brown University, Tufts University, and the Forsyth Institute.

### Availability of data and materials

The datasets used and/or analysed during the current study are available from the corresponding author on reasonable request. The datasets analyzed for this manuscript will also be available through the NCBI under the BioProject accession no.: PRJNA421501 accompanying the publication of Chung et al. (2019). R scripts utilized to perform the simulation studies and to analyze the pancreatic cancer dataset are available via the GitHub directory provided in the reference section, Meier (2019). An example script showing the analysis of a simple pseudo-dataset is also available in the same directory.

## Author’s contributions

RM developed the methodology, conducted the statistical analyses and wrote the manuscript. RM managed the acquisition and processing of the reference data sets and edited the manuscript JAT provided advice and guidance on the statistical methodology and edited the manuscript. MC, NZ and KTK assisted in the writing of the manuscript and interpretation of the study findings. DSM is the principal investigator of the pancreatic microbiome study and assisted in the interpretation and drafting of the manuscript. DCK helped conception of the methodology, supervised the implementation, and edited the manuscript. All authors read and approved the final version of the manuscript.

## Acknowledgement

We would like to extend our gratitude to Dr. Dong Pei, Lisa Neums, Stefan Graw, Qing Xia, and Duncan Rotich of the Department of Biostatistics & Data Science at the University of Kansas Medical Center for their constructive feedback on the methodology.

