## Additional File 1 for "A Bayesian framework for identifying consistent patterns of microbial abundance between body sites"

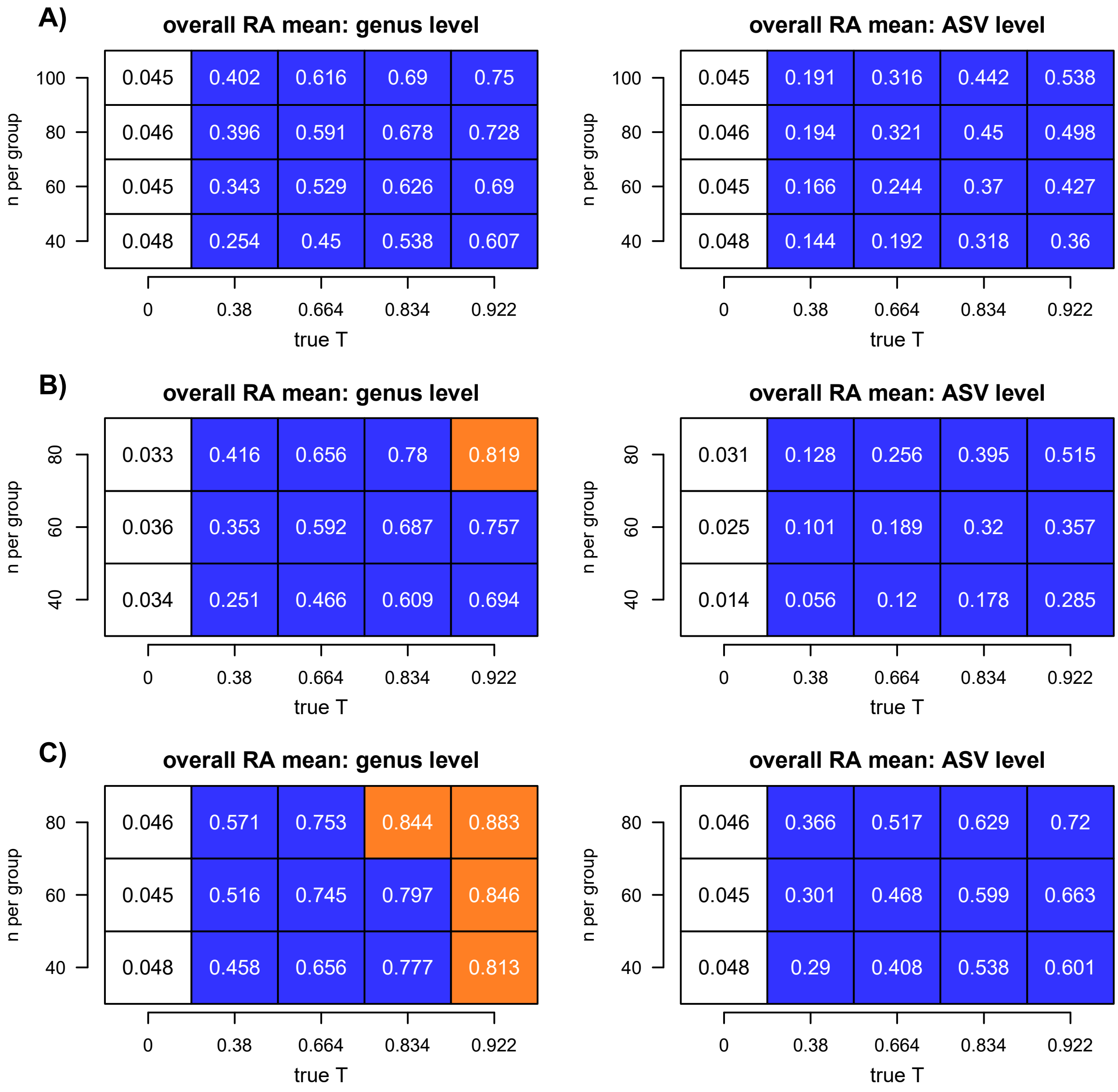


**Supplementary Figure 1:** Additional results of the simulation study for $\mu$. A) depicts the case when calibrating the lower bound of the credible interval of $T_\theta$ for a type 1 error rate of 0.05; B) depicts the case when allowing none of the strata to contain exclusively zero valued relative abundances; C) depicts the case when both conditions from A and B are met simultaneously. In part A, $H_0$ was rejected if $Pr(T_\theta|\mathbf{Y} < 0) < 0.05$. In part B and C, $H_0$ was rejected if $Pr(T_\theta|\mathbf{Y} < 0) < q$, where $q$ was adjusted for calibration. Power plots are displayed for testing PASTA of $\mu$ with $t_c = 0$ at both ASV and genus level. The term "n per group" refers to the number of samples available in each of the eight sub-group combinations resulting from two body sites and four different levels of disease status. Type 1 error rates are displayed in white colored boxes with black fonts. Power values less than 0.8 are colored blue, values larger than 0.9 are colored red and values between 0.8 and 0.9 are colored orange. While calibration does improve the power compared to the original simulations, restricting sparseness in strata leads to an even stronger improvement in performance.
