## Additional File 2 for "A Bayesian framework for identifying consistent patterns of microbial abundance between body sites"

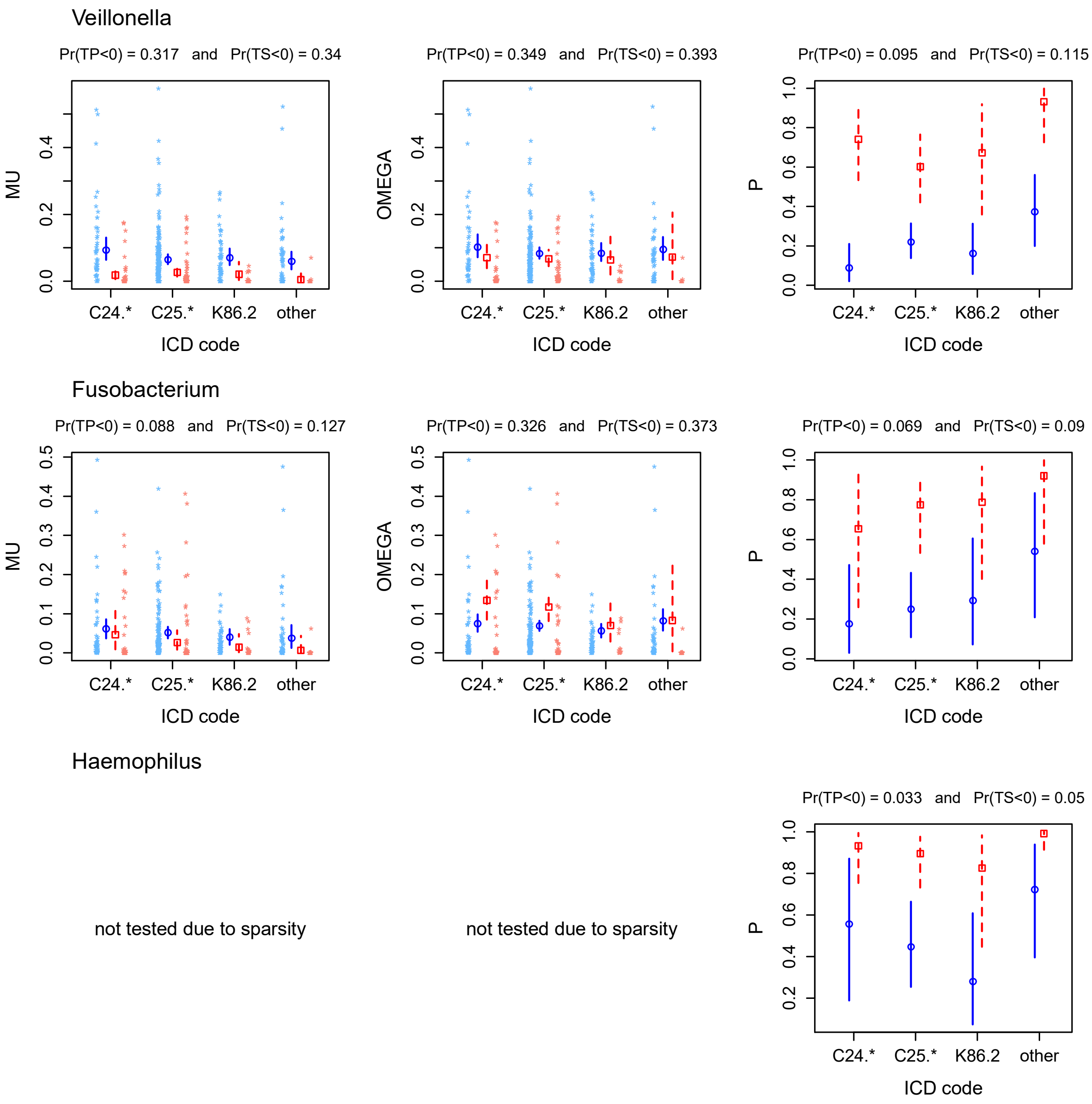


**Supplementary Figure 2:** Plots of parameter estimates within strata when testing for PASTA between gut and mouth on the genus level. Only OTUs with at least marginal significance are displayed. Each row displays the results of an OTU for the three main population parameters of interest. PASTA test results are summarized above each plot. "TP" is $T_\theta$ when utilizing Pearson correlation and "TS" is $T_\theta$ when utilizing Spearman correlation. Within each plot, circles and squares represent the posterior mean, while vertical lines represent 95% credible intervals. Body site is color coded in red and blue. For $\mu$ and $\omega$, relative abundance values are plotted next to the credible intervals.
